# Secondary metabolites and their impact on symbiotic interactions in the ambrosia fungus *Geosmithia eupagioceri*

**DOI:** 10.1101/2024.07.15.603585

**Authors:** Miroslav Kolařík, Eva Stodůlková, Soňa Kajzrová, Jaroslav Semerád, Jan Hubert, Marek Kuzma, Miroslav Šulc, Ivana Císařová, Andrej Jašica, Jan-Peer Wennrich, Jiří Hulcr, Miroslav Flieger

## Abstract

Ambrosia fungi colonize freshly dead trees, sequester nutrients, and serve as nutritional source for ambrosia beetles in exchange for dispersal. A key aspect of this symbiosis is the ability of fungi to colonize and dominate the wood around the beetle tunnels, forming a monospecific nutritional mycelium in the beetle gallery. Hypotheses for these dynamics include active beetle management, fungal inoculation priority, and the fungus’s chemical ecology facilitating resource capture and competition. The ecological role of allelochemicals produced by ambrosia fungi is unknown, although they may suppress microbes while being harmless to beetles, which has potential medical or food technology applications. This study presents a comprehensive analysis of secondary metabolites from the ambrosia fungus *Geosmithia eupagioceri* (Ascomycota: Hypocreales). Eight extracellular compounds were identified *in vitro*: 5-hydroxymethyl-2-furancarboxylic acid, 4-hydroxybenzoic acid, 2,3-dihydroxybenzoic acid, 3,4-dihydroxybenzoic acid, 4-hydroxyphenylacetic acid (4-HPA), 4-HPA methyl ester, tyrosol, and thymine. Most compounds show cross-taxon activity, suppressing the growth of bacteria, fungi, a nematode, and a mite. We have shown that often overlooked chemically simple compounds may have activities leading to increased fitness of beetle hosts, including previously unconsidered activity against mites and nematodes. For the first time, we point out that these compounds also have the previously unconsidered potential to modulate the physiology of their producer (by inducing symbiotic morphology by quorum sensing mechanisms), the beetle host and associated microbes through synergism. Furthermore, we have shown that the ambrosia fungi have biotechnological potential in the search for growth suppressors of microorganisms and invertebrates, not toxic to humans.

**IMPORTANCE:** Bark and ambrosia beetles and their microbial symbionts play crucial roles in forest ecosystems by aiding in the decomposition of dead trees, nutrient cycling, and habitat creation. However, they can cause extensive damage to both natural and planted forests by killing trees. Our study has led to a fundamental shift in the understanding of interactions between beetle symbiotic fungi and the environment, mediated by secondary metabolites. Newly, we show that these substances can not only be antimicrobial but also suppress the growth of mites, nematodes, but also can modulate the physiology of the producer fungus and potentially the host beetle and associated microbes. Our study, although conducted on a relatively artificial system with the need for validation on other lineages of ambrosia fungi, suggests entirely new research directions in the understanding of bark beetle holobiont and ambrosia beetles.

## INTRODUCTION

Plant matter primarily comprises energy sources that are not readily accessible to herbivores. As a result, certain insects such as ants, beetles, termites, and wood wasps have developed symbiotic associations with fungi to indirectly use plant substrates. This has typically been achieved by evolving dispersal benefits and culture purification benefits to the fungus, while becoming nutritionally dependent on the fungal provisioning (1). While often likened to human agriculture (2), these insect fungus associations could also be understood as evolutionary dependence of the vector on fungal metabolic capabilities, and the extent to which either beetle activity or fungal metabolism drive the ecology and evolution of the symbiosis is a subject of active research (2-4).

Among the “fungal crops”, ambrosia fungi associated with ambrosia beetles (Coleoptera: Scolytinae and, Platypodinae) include the oldest, most widespread, and most species diverse groups. The ability to feed beetle vectors and receive dispersal benefit from them evolved in at least ten groups, of which the most speciose and best studied are the ophiostomatoid ambrosia fungi (Ascomycota: Ophiostomatales) (5), closely followed by Microascales species (Ascomycota: Microascales) (6). However, current research continues to discover an ambrosial lifestyle in many groups where it was previously unknown. These includes the recently discovered basidiomycete *Irpex* spp. (Basidiomycota: Polyporales) and *Entomocorticium* (Basidiomycota: Penioporales) (7), *Fusarium* (Ascomycota: Hypocreales) (8) and *Geosmithia* (Ascomycota: Hypocreales) (9, 10).

Beetle-associated ambrosia fungi typically dominate colonized wood and beetle tunnels after inoculation. Colonization is rapid, but can be followed by a rapid decline, after which the tunnel is frequently overcome by bacteria, fungi, yeasts and invertebrates that take advantage of the nutrition-rich and secluded environment. The mechanism underlying this ecological dynamic remains unclear. One possible scenario is the priority effect, in which the early inoculation of the ambrosia fungus by the beetle enables it to capture the majority of the substrate, and only later it is outcompeted by slower-growing but more competitive wood decay and other fungi (11). Another scenario frequently mentioned in literature, is an active management of the “garden” by the beetle; while *in vitro* observations show that the beetle family interacts with the fungal conidia, it is not yet clear how this would translate into the ambient wood, where most of the fungal mass resides (12).

An additional hypothesis posits that symbiotic prokaryotes may produce chemicals to suppress fungal and bacterial competitors and invertebrate grazers, and thus confer fitness benefits to beetle inhabitants (13). Evidence exists that the ambrosia fungi themselves are likely responsible for much of the active competition with other fungi and possibly prokaryotes, having inherited a rich array of secondary chemicals that are produced into the wood during the colonization phase (14). For example, ethanol is produced by some *Ambrosiella* (15), and *Fusarium ambrosium* can synthesize naphthoquinones with antibacterial and antifungal activity (16). Secondary metabolites have not been studied in ophiostomatoid ambrosia fungi, surprisingly, the genome of the unusually pathogenic species *Harringtonia* (previously *Raffaelea*) *lauricola* has substantial potential (17). It is a tradition in the ambrosia symbiosis literature to speculate on the symbiotic implications of these products; however, this remains to be tested. Phylogeny-informed comparative studies are needed to determine whether these products are adaptations to symbiosis, an adaptation to the ecological context of the fungus, or more fundamental features of fungal metabolism that predate the evolution of symbiosis. Two comparative studies of ambrosia fungus metabolism and catabolism concluded that the vast majority of ambrosia fungus metabolism is simply inherited from their free-living ancestors, and no convergent symbiotic metabolome needs to be invoked (18).

This study aimed to document the production and biological activity of secondary metabolites in *Geosmithia eupagioceri* M. Kolarik. *Geosmithia eupagioceri* is the principal mutualist of the ambrosia beetle *Eupagiocerus dentipes* Blandford. The beetle is known from Central America and Mexico, and feeds on woody vines (19, 20). In addition to a comprehensive chemical profiling of the fungal products, we also investigate how these substances influence prokaryotes, fungi, nematodes, mites and morphology of this producer. Although we use laboratory models, rather than organisms natively co-occurring with the fungus in its native subcortical niche in the Neotropics, the phylogenetic range of the tested organisms allowed for a certain degree of generalization of the assays.

## RESULTS

Our findings suggested that *G. eupagioceri* does not produce structurally complex or novel compounds. Secondary metabolites have relatively simple structures, but their biological activity affects a broad spectrum of organisms, inhibiting bacteria, nematodes, and mites.

We identified eight compounds produced *in vitro*: 5-hydroxymethyl-2-furancarboxylic acid (**1**), 4-hydroxybenzoic acid (**2**), 2,3-dihydroxybenzoic acid (**3**), 3,4-dihydroxybenzoic acid (**4**), 4-hydroxyphenylacetic acid (**5**), 4-HPA methyl ester (**6**), tyrosol (**7**), and thymine (**8**). The compounds were structurally determined using HRMS (all compounds), NMR (all compounds except for **4**, **7**, **8**), HPLC UV-VIS by co-elution with standard compounds (**1**-**5**, **7**, **8**), and X-ray analysis (**7** and **8**) (Figure 2). Details regarding compound identification are provided in the Supplementary Material. The quantities of the dominant compounds in the fermentation broth reached 13.30 mg/L for **1**, 1.92 mg/L for **4**, and 1.21 mg/L for **8** after 7 days (Figure 3). The absolute quantities are less relevant as it is difficult to translate them from *in vitro* to *in vivo* contexts, but they show which compounds were abundant relative to one another.

**Figure 1.**
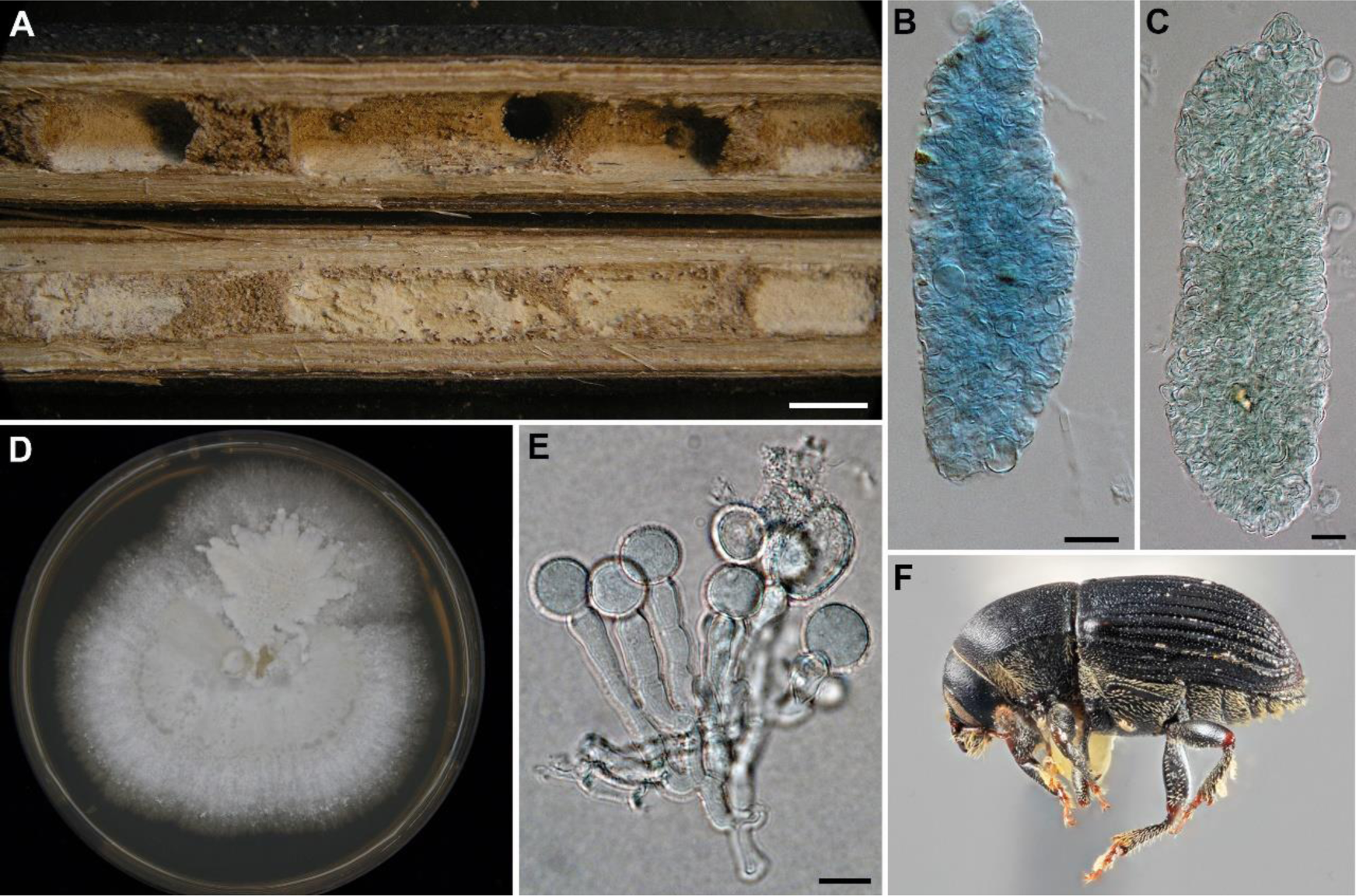
Ambrosia beetle *Eupagiocerus dentipes* and its ambrosia fungus *Geosmithia eupagioceri*. A. Gallery in *Paullinia rossii* with cream coloured ambrosia layer. B. C. Fecal pellets consisting collapsed *G. eupagioceri* hyphae and conidia, stained by cotton blue. D. Colony of *G. eupagioceri* on MEA. E. Conidiophore of *G. eupagioceri*. F. Female of *Eupagiocerus dentipes*. Scale bar. A – 5 mm, B, C, E – 20 µm.

**Figure 2.**
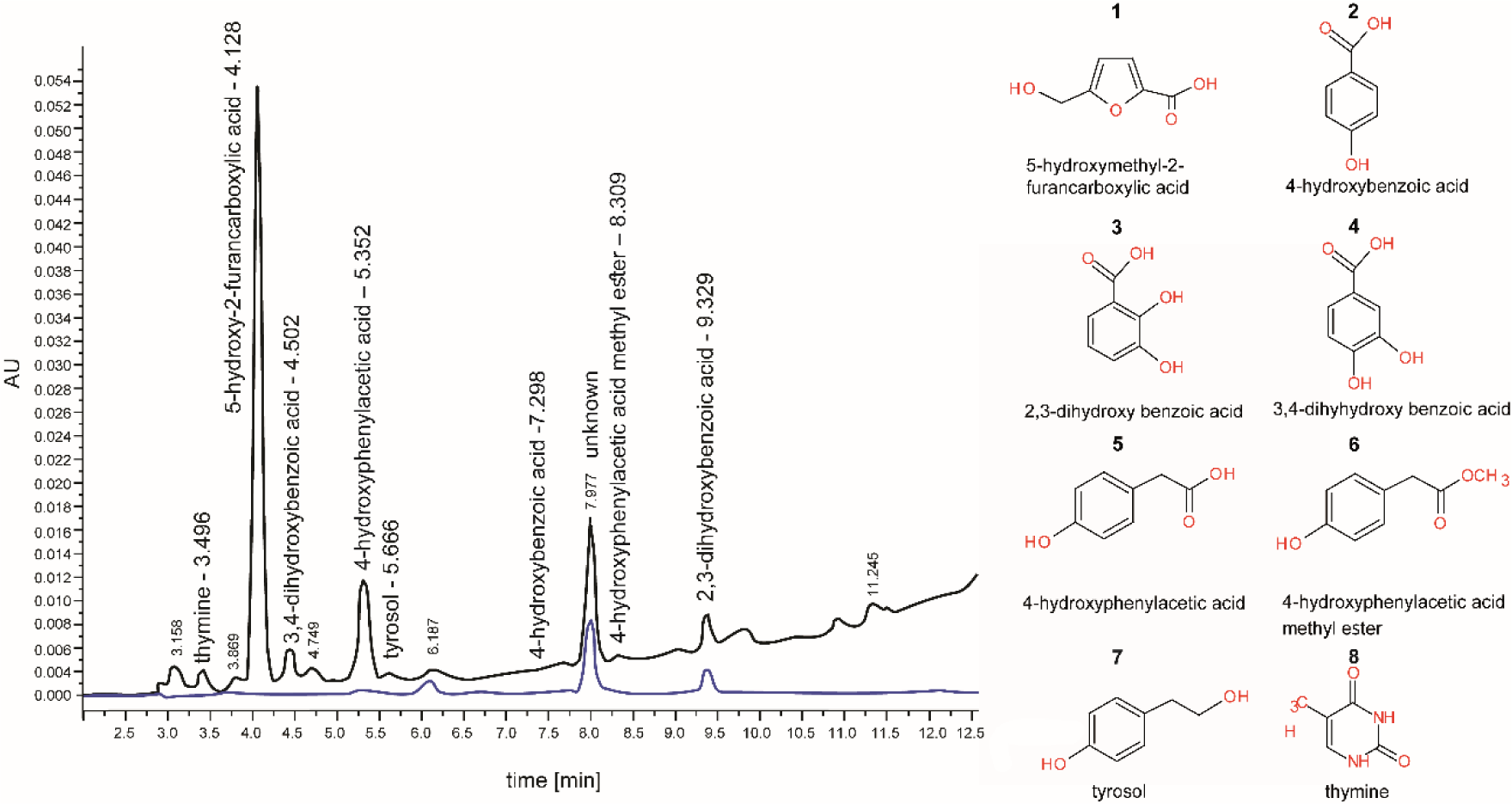
HPLC analysis of acidic ethyl acetate extract from the fermentation broth of *G. eupagioceri* (7 d). UV detection at 260 nm (black) and 340 nm (blue) is presented. Structures of the identified compounds are presented.

**Figure 3.**
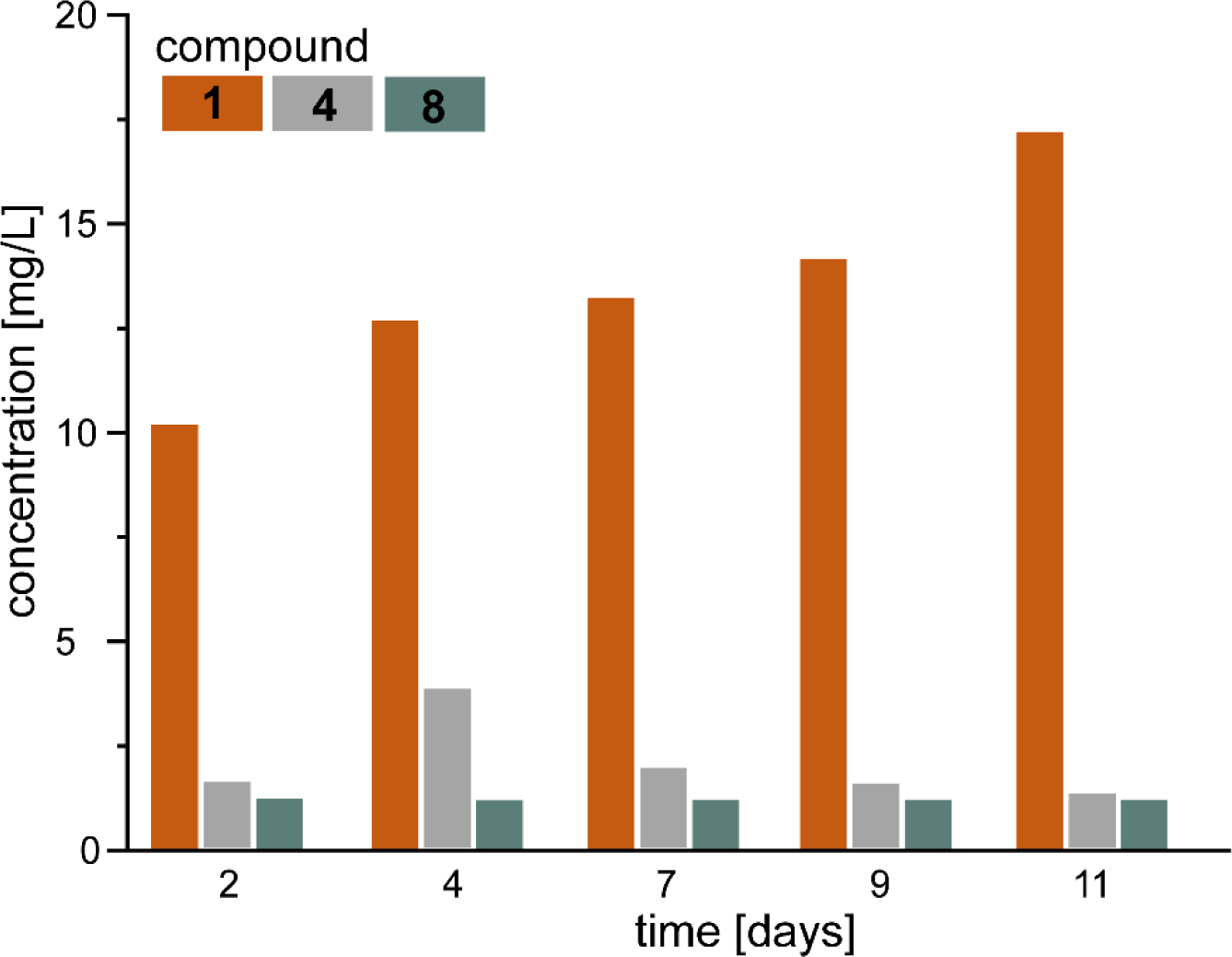
Production of the three extracellular metabolites produced in the highest concentration: **1** (5-hydroxymethyl-2-furancarboxylic), **4**(3,4 dihydroxybenzoic acid) and **8** (thymine) by *G. eupagioceri*.

Seven main compounds (all except **6**) were tested for their ability to the inhibit growth of the model fungi and bacteria. Concerning the tested fungi, **4**, **7** and **8**, had the minimum inhibitory concentration (MIC) value of 8000 μg/mL, or the MIC value was not reached. **1** had an MIC value of 8000 μg/mL for most of the strains with the exception of *Penicillium* (MIC 4000 μg/mL). **2**, **3** and **5** showed species-specific activity against most fungal species with MIC values 1000-8000 μg/mL or the MIC values were not reached. Only **1** for *Penicillium*, **2** for *G. rufescens* and *Metarhizium* (MIC 2000 μg/mL) and **5** for *G. rufescens* (MIC 4000 μg/mL), *Saccharomyces* and *Mucor* (1000 μg/mL) demonstrated inhibitory capabilities against fungi at ecologically relevant concentrations, that is, that would not concomitantly inhibit their producer, *G. eupagioceri*. In the case of bacteria, **7** and **8** had MIC value 8000 μg/mL or the MIC value was not reached. The other compounds inhibited both bacteria at concentrations of 250-2000 μg/mL. These compounds, with the exception of **3** against *Kocuria* (MIC 1000 μg/mL), inhibited bacteria at concentrations lower than those inhibiting *G. eupagioceri* and could be considered ecologically significant (Table 1).

**Table 1.** Minimal Inhibitory Concentration (MIC) of the seven studied compounds for filamentous fungi, yeast, and bacteria. Compounds were tested in the concentration range 8000, 4000, 2000, 1000, 500, 250, 125 and 63 ug/ml. Values below the MIC of *G. eupagioceri* are highlighted in bold.

| Microbe | MIC (µg/mL) |  |  |  |  |  |  |  |  |  |
| --- | --- | --- | --- | --- | --- | --- | --- | --- | --- | --- |
|  | **1**<br>(HMFC<br>A) | **2** (4-HBA) | **3** (2,3-DHBA) | **4** (3,4-DHBA) | **5** (4-HPA) | **7**<br>(tyros<br>ol) | **8**<br>(thym<br>ine) | chlora<br>mpheni<br>col | amphot<br>ericin B | cyclohe<br>ximide |
| *G. eupagiocer*<br>*i* | 8000 | 4000 | 1000 | 8000 | 8000 | 8000 | >800<br>0 | n.a. | <0.13 | <0.50 |
| *G. rufescens* | 8000 | **2000** | 2000 | 8000 | **4000** | 8000 | >800<br>0 | n.a. | <0.13 | <0.50 |
| *Metarhiziu*<br>*m*<br>*anisopliae* | 8000 | **2000** | 2000 | 8000 | >800<br>0 | 8000 | >800<br>0 | n.a. | 2.083 | >66.66 |
| *Mucor*<br>*plumbeus* | 8000 | 4000 | >8000 | 8000 | **1000** | >800<br>0 | >800<br>0 | n.a. | <0.13 | >66.66 |
| *Beauveria*<br>*bassiana* | 8000 | >800<br>0 | 1000 | 8000 | 8000 | 8000 | >800<br>0 | n.a. | 2.083 | >66.66 |
| *Penicillinu*<br>*m*<br>*decumbens* | **4000** | 4000 | 2000 | 8000 | >800<br>0 | 8000 | >800<br>0 | n.a. | 16.6 | >66.66 |
| *Saccharom*<br>*yces*<br>*cerevisiae* | 8000 | 4000 | 4000 | 8000 | **1000** | 8000 | >800<br>0 | n.a. | <0.50 | <0.50 |
| *Escherichi*<br>*a coli* | **1000** | **500** | **500** | **2000** | **250** | 8000 | >800<br>0 | 1.04 | n.a. | n.a. |
| *Kocuria*<br>*rhizophila* | **2000** | **1000** | 1000 | **1000** | **250** | 8000 | >800<br>0 | <1.04 | n.a. | n.a. |

**Table 1.** Minimal Inhibitory Concentration (MIC) of the seven studied compounds for filamentous fungi, yeast, and bacteria. Compounds were tested in the concentration range 8000, 4000, 2000, 1000,
| Microbe | MIC (µg/mL) |  |  |  |  |  |  |  |  |  |
| --- | --- | --- | --- | --- | --- | --- | --- | --- | --- | --- |
|  | **1**<br>(HMFC<br>A) | **2** (4-HBA) | **3** (2,3-DHBA) | **4** (3,4-DHBA) | **5** (4-HPA) | **7**<br>(tyros<br>ol) | **8**<br>(thym<br>ine) | chlora<br>mpheni<br>col | amphot<br>ericin B | cyclohe<br>ximide |
| *G. eupagioceri* | 8000 | 4000 | 1000 | 8000 | 8000 | 8000 | >800<br>0 | n.a. | <0.13 | <0.50 |
| *G. rufescens* | 8000 | **2000** | 2000 | 8000 | **4000** | 8000 | >800<br>0 | n.a. | <0.13 | <0.50 |
| *Metarhizium anisopliae* | 8000 | **2000** | 2000 | 8000 | >800<br>0 | 8000 | >800<br>0 | n.a. | 2.083 | >66.66 |
| *Mucor plumbeus* | 8000 | 4000 | >8000 | 8000 | **1000** | >800<br>0 | >800<br>0 | n.a. | <0.13 | >66.66 |
| *Beauveria bassiana* | 8000 | >800<br>0 | 1000 | 8000 | 8000 | 8000 | >800<br>0 | n.a. | 2.083 | >66.66 |
| *Penicillium decumbens* | **4000** | 4000 | 2000 | 8000 | >800<br>0 | 8000 | >800<br>0 | n.a. | 16.6 | >66.66 |
| *Saccharomyces cerevisiae* | 8000 | 4000 | 4000 | 8000 | **1000** | 8000 | >800<br>0 | n.a. | <0.50 | <0.50 |
| *Escherichia coli* | **1000** | **500** | **500** | **2000** | **250** | 8000 | >800<br>0 | 1.04 | n.a. | n.a. |
| *Kocuria rhizophila* | **2000** | **1000** | 1000 | **1000** | **250** | 8000 | >800<br>0 | <1.04 | n.a. | n.a. |

We further studied the effect of **1**, **2**, **4**, **5**, **7** and **8** (selected to represent the breadth of the excreted chemistry) on the *G. eupagioceri* morphology. In all cases, including the control (water or ethanol), the conidia germinated into hyphae. The control mycelium exposed to **2**, **5**, and **7** as well as the mycelium exposed to other substances at a concentration of 500 µg/ml, did not sporulate and formed branched hyphae, 2-4 µm wide, without swollen cells. In presence of **1** (4000-2000 ug/ml) and **4** (8000-1000 ug/µl) the apparent presence of monillioid hyphae with swollen hyphal elements, 3.5-8.0 µm wide, was observed. The most pronounced effect was observed with **8** (8000-500 µg/ml), which induced the frequent presence of monillioid hyphae with swollen cells, conidiophores, and conidia (Figure S2).

None of the tested compounds appeared to have synergistic effects on bacteria; the effect was additive.

Three substances produced by *G. eupagioceri* at the highest concentrations, **1**, **7**, and **4**, were screened for their effects on nematode and mite model. Statistically significant nematicidal activity was observed at the highest concentrations: 400 µg/ml for **1** (Kruskal-Wallis, P < 0.05), 400 (P < 0.01) and 200 µg/ml (P < 0.05) for **7**, and 400 (P < 0.01) and 200 µg/ml (P < 0.01) for 3,4-DHBCA (Figure 3).

The three tested compounds decreased the population growth of the mite *Tyrophagus putrescentiae* (Kruskal-Wallis H(chi^2^) = 77.7, P < 0.001). Except for **7** at 0.01 mg/ml, all tested compounds showed a significant decrease in mite density (Mann-Whitney pairwise test, Bonferroni corrected P < 0.05). The highest toxic effect was observed with **7** at 1 mg/ml, reducing the mite population density by 8-fold compared with the control (Figure 4). In comparison to the control, **4** at 1 mg/ml and **1** at 1 mg/ml decreased the mite population density by 6 and 4.5 times, respectively.

**Figure 4.**
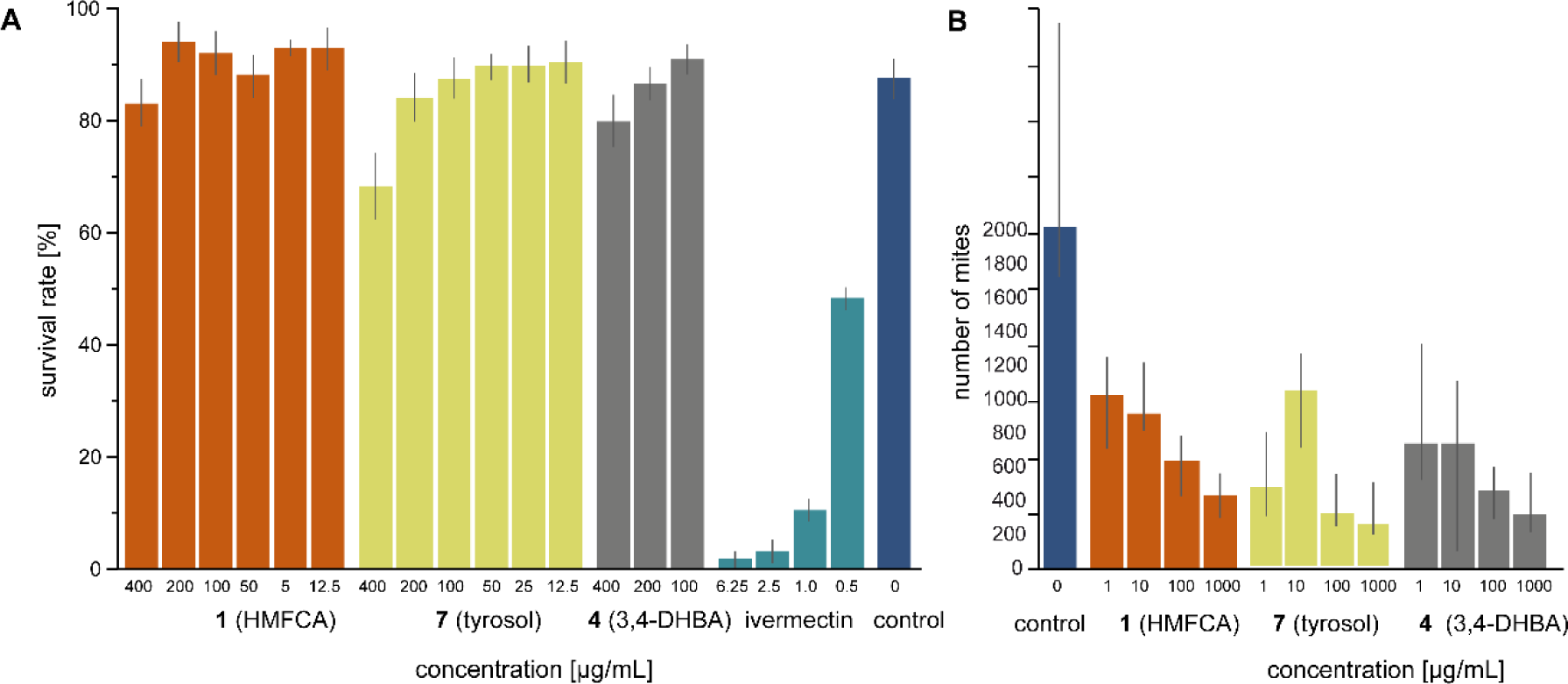
A. Nematicidal activities of **1**, **4, 7** and positive control (ivermectin) and negative control against *Caenorhabditis elegans* in the microtiter plate assay. **B**. The population density of *Tyrophagus putrescentiae* after 21 days of cultivation from 10 individuals on control and tested compounds treated diets. In graph B, the columns are medians and bars represent interquartile range. Note that the assay reports survival in nematodes and fecundity in mites, given the different biology of the two models.

## DISCUSSION

*G. eupagioceri* produced eight compounds. All are structurally simple, and already known as fungal metabolites: **1** [5-hydroxymethyl-2-furancarboxylic acid, Sumiki’s acid, 5-(hydroxymethyl)-2-furoic acid, HMFCA] (21-25), **2** (4-hydroxybenzoic acid, *p*-hydroxybenzoid acid, 4-HBA) (26), **3** (2,3-dihydroxybenzoic acid, 2,3-DHBA) (27), **4** (3,4-dihydroxybenzoic acid, protocatechuic acid, 3,4-DHBA) (28, 29), **5** (4-hydroxyphenylacetic acid, *p*-hydroxyphenylacetic acid, 4-HPA) (Ohtani, Fujioka et al. 2011), **6** (4-hydroxyphenylacetic acid methyl ester) (30), **7** (tyrosol, 2-(4-hydroxyphenyl)-ethanol)) (21, 31-34), and **8** (thymine) (35). All compounds originated from three biosynthetic pathways. Benzoic acid derivatives, **5** and **7** are part of the phenylpropanoid pathway, and originates from phenylalanine or tyrosine, both of which are derived from shikimate (36, 37). **1** is a furan compound produced from hexoses through several enzymatic steps (38). **8** is a pyrimidine nucleobases that can be obtained by fungi through *de novo* biosynthesis or salvage (recycling) (39). As *G. eupagioceri* can grow on basal Czapek-Dox medium without added nucleosides (9), we assume that it is capable of *de novo* synthesis.

### Secondary metabolites – allelopathic abilities

The biological activity of the identified compounds can be summarized as follows: all except **7** and **8** suppress the growth of bacteria, few of them (**1**, **2**, **5**) suppress the growth of fungi and yeasts, and the three that are produced in the highest concentration suppress mites and mites, within an ecologically realistic range of concentrations (i.e. concentrations lower than those that inhibit the producer. This suggests that these secondary metabolites may have antibacterial and anti-invertebrate functions in nature, and that their effects against fungi are marginal. They are unlikely to evolve a symbiotic relationship with a beetle if they produce an insecticidal compound; which is consistent with the fact that the metabolites are not toxic to animals, including mammals, even at high doses, and are already utilized as additives and preservatives in the food industry (40-43).

Our findings regarding the biological activities are consistent with those of previous studies. We did not test the compounds on organisms sympatric with the fungus in its original habitats. Marginal antifungal activity has been shown for **1** (23, 24), **3** (44), **5** (45) and **7** (46, 47). Although the model organisms used here span great phylogenetic breadth, we refrain from overgeneralizing our results, because these compounds may have different effect on different species of bacteria and invertebrates (see below). Species-specific antibacterial activity has already been reported, with the MIC values corresponding to our results. For instance, **1** did not show any inhibitory activity against several bacteria, including *E. coli*, at the highest tested concentrations (256 µg/ml, Quang et al. (2022). However, other bacterie, such as *Enterococcus faecalis* (23), *Ralstonia solanacearum* (31 µg/ml), *Agrobacterium tumefaciens* (31 µg/ml), and *Erwinia carotovora* (16 µg/ml) (22), were highly sensitive. Antibacterial activity was reported for **7** (46, 48) and **3** (George et al. 2011, MIC not determined). Further for **4** with MIC 8-64 μg/ml (40), 2.7 mg/ml (49), 1 and ˃1 mg/ml (50). In **2** the reported values were IC50 160 μg/ml (51) and MIC 1 and ˃1 mg/ml, (50). For **5** the MIC was 1 and ˃1 mg/ml, (50) and LD90 1 mg/mL (52). Alteration of bacterial growth stems not only from inhibition but also from the prevention of biofilm formation. This phenomenon has been documented with various phenolic compounds, including **3** (53) and its precursor, **7** (48). Moreover, previous studies have revealed antiprotozoal activity in **3** (active at 2.5 mg/ml, (54)), as well as antiviral activity in **1** (24) and 3,4-DHB (42).

Our study showed the significant nematocidal effects of **1** and **4**. These findings, including the concentrations that induce inhibition, are in line with those of numerous studies. **1** effectively combats the pine wood nematode *Bursaphelenchus xylophilus* and *Caenorhabditis elegans* (approximately 20% mortality after 6 days, 284 µg/ml), but showed no activity against *Pratylenchus penetrans* (55). It also significantly reduced the viability of the root knot nematode *Meloidogyne incognita* at concentrations of 400, 200, and 100 µg/ml, and inhibited egg hatching at 200 µg/mL. Consequently, it is considered a primary tool of its producer, the nematophagous fungus *Drechmeria coniospora*, in nematode-killing (25). **4** displays nematicidal activity against *Meloidogyne incognita* with an MIC below 125 µg/ml (56). Lastly, 4-HPA exhibited activity (22% inhibition at a concentration of 456 μg/ml) against *Pratylenchus penetrans* and *Bursaphelenchus xylophilus* (57).

Our study further revealed significant acaricidal properties of the three tested substances. The toxic effects of these compounds were previously undocumented, but are known in related substances, such as benzoic acid (58), 6-[(Z)-10-heptadecenyl]-2-hydroxybenzoic acid (59), and benzyl benzoate—an established, and widely-used commercial acaricide (60). The mode of action of these compounds is not completely understood; however, mites are dependent on bacterial symbionts and the observed compounds show strong antibacterial effects (61, 62).

### Secondary metabolites – quorum sensing

Numerous fungi utilize small molecules as quorum sensing (QS) signals, for intraspecies as well as interspecies or interkingdom communication. The capacity to influence bacterial metabolism and sensitize them to antibiotics has been observed for benzoic acid, **2** (63, 64), **8** (4); **4** (65), **3** (66), and **7** (47, 67). Additional quorum sensing activity is recognized in **2** and **4**, which can influence metabolism, pathogenicity, and antifungal compound production in various bacteria (68, 69). In our study, we did not find synergistic antibiotic activity; however, the exact mode of action of these compounds remains unclear, and interference with bacterial ecology such as the prevention of adhesion or quorum sensing, rather than antibiosis, remains a possibility.

The capacity of the identified substances to modulate hyphal morphology was noteworthy. Ambrosia fungi spend most of their life cycle as filamentous mycelium in wood, but critical parts of their ecology are concentrated in the ambrosia layer with swollen cells and conidia on which the insect symbiont feeds, and the monilioid mycelium inside the mycangia, an organ for purification of the fungal culture and its maintenance during dispersal. The transition from yeast to hyphae involves intricate changes in gene expression and signalling pathways. Remarkably, three identified compounds, **2** in *Candida albicans* (68), **5** in *Ophiostoma ulmi* (70), and **7** in *C. albicans* (71, 72), impact the yeast-to-hypha transition, leading to diminished hyphal formation. Moreover, **3** stimulates the germination of *Colletotrichum musae* (73). In our study, we did not observe any effect on the transition between the yeast and hyphal phases. However, with several substances (**1**, **4** and **8**), we observed an influence on the formation of monillioid hyphae, characterized by swollen cells, as well as the presence of conidiophores and conidia.

It is also possible that, rather than having an ecological role, these high-concentration products facilitate fungus feeding and nutrient mobilization. Benzoic acid and its derivatives function as chelators, aiding the solubilization of metals and other nutrients from organic materials. This process renders the nutrients accessible to fungi, and subsequently, insect uptake. Notably, 2,3-DHB is effective in iron chelation (74), and its capability to bind iron ions is considered a contributor to blue stain formation in ophiostomatoid fungi (27). Some of these compounds have features relevant to animals. **4**, **5** and **7** exhibits strong antioxidant properties (75, 76). **4** is involved in modulating insect physiology, affects the hardening of insect cuticles in cockroaches (77) and stimulates oviposition in ladybirds (78). The effects of *G. eupagioceri* on beetles remain intriguing.

### Conclusion and future perspectives

The ecologically active extretome of the fungus can be composed of well-known, long recognized and simple chemistries. However, these are typically ignored in ecological and also bioprospecting studies that focus on drug-like and novel structures. Interestingly, these substances have occasionally been reported in ophiostomatoid fungi associated with phloem-feeding bark beetles: **7**, **3** and **5** in *Ceratocystis montium*, *Grosmannia clavigera* and *G. huntii* (27), **7** in *Catunica adiposa* (syn. *Ceratocystis adiposa*) (79), and **5** in *Ophiostoma ulmi* (70, 80). We do not know whether these substances are produced by other ambrosia fungi, such as the more widespread *Ambrosiella* or *Raffaelea*, and future exploration of ambrosia fungus secondary metabolites, and tests of their ecological significance are promising frontiers in this field of research.

Similar chemicals are increasingly reported to be produced by fungi associated with ambrosia beetles or phloem-feeding bark beetles (15, 81, 82). Some were apparently harmless to their hosts (83). Ultimately the research field will need to contrast the apparent benefits between coevolved and non-coevolved fungi and carefully assess the actual titer of these compounds in the micro-habitat of the fungal hyphae.

Varied concentrations, timing and diverse chemical interactions may give rise to emergent phenomena hitherto never considered in the ecology of ambrosia fungi. Related substances such as **2, 3, 4** and **7** are known for their nonlinear biological effects, such as synergistic activity and enhancement of the antibiotic activity of co-occurring compounds. Synergistic effects are important for drug efficacy (84), but this feature has been little studied in microbial ecology. Ophiostomatoid fungi produce a range of volatile terpenoids, a chemical class with known synergistic activities (85). Several of the identified compounds are capable of stimulating swollen cells and conidiophore formation, that is growth forms observed in ambrosial beetle galleries. Morphogenetic stimuli may originate from the fungi themselves, rather than from the beetles, and their activity may be a function of abundance or a confined space.

Our study shows that ambrosia fungi can produce substances with a range of biological activities. The advantage of the studied system is that these substances are not toxic to insects, which increases the chances of their use in human medicine. This makes ambrosia fungi, with their enormous taxonomic diversity, suitable candidates for biotechnology.

## MATERIALS AND METHODS

### Strain and cultivation

*Geosmithia eupagioceri* CCF 6377 was isolated from the ambrosia beetle *Eupagiocerus dentipes*, collected in Belize, Las Cuevas Research Station, January 28, 2019, as no. 18063 by You Li and J. Hulcr under Scientific collection/Research permit wildlife protection act no. 14/2000 (no. QL/2/2/18(35)) issued by the Ministry of Agriculture, Fisheries, Forestry, Environment and Sustainable Development of Belize. The strain was identified using the morphology and ITS rDNA sequencing using the methods of Kolařík et al. (86) and the sequence is deposited in NCBI Genbank under to number PP923662. This strain was deposited in the Culture Collection of Fungi (CCF, Department of Botany, Charles University, Prague, Czech R.). The strain was cultivated on liquid MEA (MEA; 20 g malt extract (Oxoid), 20 g glucose, 1 g peptone (Difco), 1000 ml distilled water, pH 6.5) medium for 14 days.

### Extraction, isolation and metabolite identification

Metabolites were studied in cultivation broth. The chemicals were extracted with ethyl acetate and acidic ethyl acetate and the combined extracts were concentrated under reduced pressure. The crude extract was loaded onto a 5g C18 solid phase extraction (SPE) cartridge. Compounds were eluted with step gradient H_2_O/MeOH + 0.1% TFA and obtained fractions were further purified by semi preparative RP HPLC (Gemini, isocratic elution, H_2_O/MeOH + 0.1% TFA, effluent monitored at 260 nm (87). Qualitative analysis of the separated compounds was performed using a high-resolution mass spectrometer (Agilent 6546, qTOF) equipped with an electrospray ion source operating in positive and negative modes. A Poroshell 120 EC-C18 column (2.7 µm, 3 ×100 mm) and gradient elution using water with the addition of formic acid (A) and methanol (B) were used to separate individual substances. The method consisted of an initial isocratic phase for 0.5 min (30% B) followed by gradient elution from 30% B to 100% B over 15 min. After 3 min of 100% B, default method conditions were reset, and the system was equilibrated for 5 min before the next injection. The flow rate was set to 0.4 ml/min and 40 °C. The settings of MS operating in negative ionization mode were as follows: sheath gas temperature and flow, 400 and 12 L/min; drying gas temperature and flow, 250°C and 8 L/min; nebulizer pressure 35 psi; capillary voltage, -3500 V; fragmentor, skimmer, and Oct 1 RF Vpp, 140, 65, and 750 V; mass range of 50-1700 m/z; collision energy 0, 20, and 40 eV; acquisition rate of 3 spectra/s. Acquired data were analyzed using Agilent MassHunter Qualitative Analysis software (version 10.0). Detailed HRMS data for the identified compounds are provided in Supplementary material 1.

Seven of the eight identified compounds, **1** (No. H40807, 99%), **2** (No. PHR1048, 99%), **3** (No. 126209, 99%), **4** (No. 37580, ≥97.0%), **5** (No. H50004, 98%), **7** (No. 188255, 98%), **8** (No. T0376, ≥99.0%) (all from Sigma-Aldrich) were used as HPLC, HRMS, and NMR standards and for all bioassays. Because some of these compounds might be present in trace amounts in the malt extract itself, we also investigated the pure liquid 2% MEA as a negative control.

### Extracellular metabolite quantification by HPLC

Three main components were further quantified during the time course of liquid cultivation, based on the calibration curves. *Calibration.* Standard solutions of each tested compound (**1**, **4**, and **8**) were prepared in methanol at final concentrations of 15.5, 31, 62.5, and 125 mg mL^-1^. Calibration graphs were constructed by plotting the integrated peak areas of the individual compounds versus the concentration. The following linear regression equation and correlation coefficient were obtained: **1**, y = 8041.7x – 41004, R² = 0.999; **4** y = 6844.6x – 34559, R² = 0.9989; **8**, y = 30451x – 136934, R² = 0.9991.

*NMR analysis.* The structure elucidation using NMR was carried out on a Bruker Advance III 700 MHz spectrometer (700.13 MHz for ^1^H, 176.05 MHz for ^13^C) and Bruker Advance III 600 MHz spectrometer (600.23 MHz for ^1^H, 150.93 MHz for ^13^C) in acetone-*d_6_* and CD_3_OD at 20°C. The spectra were referenced by the residual signal of the solvent (for acetone-*d_6_* δ_H_ 2.060 ppm, δ_C_ 29.86 ppm, CD_3_OD δ_H_ 3.305 ppm, δ_C_ 49.04 ppm). Pulse sequences provided by the manufacturer of the spectrometers were used to for the NMR experiments: ^1^H NMR, ^13^C NMR, COSY, ^1^H-^13^C HSQC, and ^1^H-^13^C HMBC. The ^1^H NMR and ^13^C NMR spectra were zero-filled to fourfold data points and multiplied by the window function before Fourier transformation. A two-parameter double-exponential Lorentz-Gauss function was applied to ^1^H to improve the resolution and line broadening (1 Hz) was applied to obtain a better ^13^C signal-to-noise ratio. Chemical shifts are given in δ-scale with digital resolution justifying the reported values to three (δ_H_) or two (δ_C_) decimal places. Detailed NMR data for the identified compounds are provided in Supplementary material 1.

*X-Ray analysis.* Two of the isolated compounds were obtained in the crystal form, and their identification was conducted using X-ray crystallography. Diffraction experiments were performed on a Bruker D8 VENTURE Kappa Duo PHOTONIII with an IμS micro-focus sealed tube CuKα (λ= 1.54178 Å) at a low temperature of 120K. The structure was solved by direct methods (XT (88)) and refined by full matrix least squares based on *F*^2^ (SHELXL2018 (89)). The hydrogen atoms on carbon were fixed into idealized positions (riding model) and the temperature factor H_iso_(H) = 1.2 U_eq_(pivot atom). The hydrogen atoms on oxygen were found on the difference Fourier map and refined under the assumption of a rigid body with temperature factors H_iso_(H) = 1.2 U_eq_(pivot oxygen). The absolute configuration of the crystal is based on anomalous scattering of O and N atoms; however, the large standard deviation of the Flack parameters suggests a low reliability of the determination. X-ray crystallographic data of **7** were obtained for the first time and have been deposited with the Cambridge Crystallographic Data Centre (CCDC) under deposition number 2158213 and can be obtained free of charge from the Center via its website (www.ccdc.cam.ac.uk/getstructures). The structural characteristics are presented in Supplementary material 2.

### Antimicrobial activity and effect on phenotype

Minimum inhibitory concentrations (MIC) were determined in a serial dilution assay (concentration range 63-8000 µg/ml) on two bacterial and seven fungal species. The tested strains included the source organism (*G. eupagioceri*), and model species of gram-positive and gram-negative bacteria (*Kocuria rhizophila* CCM552, =ATCC9341; *Escherichia coli* ATCC3988), yeast *Saccharomyces cerevisiae* CCM8191 (=ATCC9763), filamentous saprophytic fungi *Geosmithia rufescens* CCF4524, *Penicillinum decumbens* CCF4423, *Mucor plumbeus* CCF2612, and entomopathogenic fungi *Beauveria bassiana* CCF4422 and *Metarhizium anisopliae* CCM2787. For all assays, bacteria were cultivated on LB broth (Sigma-Aldrich, pH 7.0), and yeasts and filamentous fungi were cultivated on yeast-malt broth YM6.3 (glucose 4 g, malt extract 10 g, yeast extract 4 g, H_2_0 1L, pH 6.3). The inoculum suspensions were prepared from liquid culture cultures (1-7-day-old, depending on the taxon) and adjusted to inoculum an density of 0.4 × 10^4^, as demonstrated by quantitative colony counts. The mixture was incubated for 24 (bacteria and yeasts) or 48 h (filamentous fungi) at 25 °C on a shaker (300 rpm). The tests were conducted in triplicate in 96-well microtiter plates and were visually evaluated. The antibiotics, amphotericin B, chloramphenicol, and cycloheximide, in concentration ranges specific for each compound, were used as quality control standards. The compounds were dissolved in ethanol (**2, 8**) or water (others). The toxicity of ethanol was considered in all measurements; thus, negative controls also contained EtOH concentrations equivalent to those present with each tested substance. MICs were defined as the lowest concentration of the test compound at which the test organism did not grow (filamentous fungi, bacteria) or the growth was partially inhibited (> 50 % inhibition) for yeasts, as recommended for each group (90). See Štěpánek et al. (91) for further details.

Because some of the identified compounds have been described to influence the phenotype of fungi, we tested the effects of **1**, **2**, **4**, **5**, **7** and **8** on the morphology of their producer, *G. eupagioceri*. These compounds were selected to cover the dominant compounds that represent the entire chemical diversity. The dimension and shape of hyphae, as well as the presence of sporulation, were monitored using light microscopy directly from the wells of the 96-well plates used in the MIC study.

We also assessed the synergistic effect of the same set of compounds on *S. cerevisiae* and both bacteria. Because of the substantial number of combinations, we tested only their 1:1 weight proportion mixture and evaluated concentrations in the range of 63-2000 µg/ml. Thus, in the case of 2000 µg/ml, it indicates the addition of 1000 µg/ml of each compound to the mixture.

### Nematicidal activity

*Caenorhabditis elegans* was inoculated monoxically on nematode agar. The medium was prepared from soy peptone 2,5 g/l, NaCl 3 g/l, and agar-agar 17 g/l, pH 7.4, and, after autoclaving, the following ingredients were added as sterile filtered solutions: cholesterol (1 mg/ml dissolved in EtOH) 1 ml, 1M CaCl_2_ 1 ml, 1M MgSO_4_ 1 ml, and 1M potassium phosphate buffer 25 ml, pH 6. 8. with living *Escherichia coli* (DSM 498) (1 ml of a suspension containing approximately 10 cells/ml, pre-inoculated for 12 h at 37 °C). Nematodes were added and incubated at 21 °C for 4–5 days. Nematodes were washed from the plates for use in experiments with M9 buffer (3 g KH_2_PO_4_, 6 g Na_2_HPO_4_, 5 g NaCl, 1 l H2O and, after autoclaving, the addition of 1 ml 1M MgSO_4_). 0.5 ml of the suspension containing 500 nematodes/ml was added to each well of the 24-well plate. Six concentrations (400, 200, 100, 50, 25, 12,5 μg/ml) of **1**, **4** and **7** were tested (total volume of 1 ml/well) in triplicate. Ivermectin (Sigma-Aldrich) was used as the control. The plates were incubated at 24 °C in the dark and nematicidal activity was determined after 18h of incubation.

### Acaridical activity

As a mite model, the astigmatid mite *Tyrophagus putrescentiae* (Schrank, 1781) culture “5L” was used in the experiment. The culture was collected by E. Zdarkova in a grain store in Bustehrad, Czechia in 1996, and propagated in the lab. The rearing medium was a stored product mite diet (SPMd): a mixture of oat flakes, wheat germ, and dried yeast (Rapeto, Bezdruzice, Czechia) (10:10:1 w/w). The rearing conditions have been described previously (92)

Compunds, **1, 4** and **7** were selected to represent the dominant compounds across the identified chemical diversity. The samples were diluted in double-distilled water to obtain concentrations of 0, 0.001, 0.1, 0.1 and 1 mg/mL. We used a modified protocol for population growth tests, originally developed to test the effect of residual pesticide activity (93). The diluted compounds (5 mL) were added to 10 g of SPMd in 50 mL plastic vials. The vials and enriched diets were lyophilized and then rehydrated for 24 h in an exicator with distilled water before 0.5 ± 0.05 g was added to 70 mL flasks (IWAKI flasks; Cat. No. 3100–025; Sterilin, Newport, UK). Ten unsexed adult mites were added to each vial. The vials were incubated at 25 ± 1 °C and 85% RH in darkness for 21 days. Ten replicates per concentration of each compound were used. The experiment was terminated by filling the vial with 50 mL of Oudemans fluid for fixation (the fluid contained 87 mL of 70% ethyl alcohol, 8 mL of glacial acetic acid, and 5 mL of glycerol). The contents of each vial were mixed to obtain a homogenous distribution of the fixed mites. For each vial, three subsamples of 1 mL were taken and mites were counted under a dissecting microscope. The number of mites in each vial was calculated as the mean from each subsample. The total number of mites per chemical concentration was than calculated. The effect of acaricidal compounds was tested using the Kruskal-Wilcoxon test. The toxic concertation/compounds were suggested those with significant decrease in the final mite population density in comparison to the control using the Mann-Whitney paired test. All calculations were performed using the PAST 4.13 (94).

## SUPPLEMENTAL MATERIAL

## DATA AVAILABILITY

X-ray crystallographic data of **7** were obtained for the first time and have been deposited with the Cambridge Crystallographic Data Centre (CCDC) under deposition number 2158213 and can be obtained free of charge from the Center via its website (www.ccdc.cam.ac.uk/getstructures). Structural characteristics are provided in Supplementary material 2. The barcode ITS rDNA sequences of the strain was deposited in the NCBI GenBank database.

## ACKNOWLEDGMENTS

We are grateful to S.M. Smith (Michigan State University) for providing the picture of *Eupagiocerus dentipes* which is distributed under a CC0 1.0 license and is available online: https://www.barkbeetles.info. Work was supported by the Southeast Asia–Europe Joint Funding Scheme (SEA–Europe Grant number JFS20ST-127, Acronym: Antiviralfun. The authors benefited from the H2020-RISE project Mycobiomics–Joining forces to exploit the mycobiota of Asia, Africa, and Europe for beneficial metabolites and potential biocontrol agents, using -OMICS techniques (No. 101008129). The authors would like to thank our collaborators in Belize for their support in obtaining the fungus: the Belize Forest Department, the Friends of Conservation and Development, and Bull Ridge Ltd.

